# Diploid Hepatocytes Resist Acetaminophen-Induced Liver Injury Through Suppressed JNK Signaling

**DOI:** 10.1101/2025.07.31.667940

**Authors:** Sierra R. Wilson, Evan R. Delgado, Frances Alencastro, Rosa L. Loewenstein, Madeleine P. Leek, Leah R. Peters, Kerollos Kamel, Patrick D. Wilkinson, Siddhi Jain, Joseph Locker, Silvia Liu, Bharat Bhushan, Andrew W. Duncan

## Abstract

**Background & Aims:** The liver contains both diploid and polyploid hepatocytes, but their functional differences remain poorly understood. Emerging evidence suggests that each ploidy state contributes to regeneration in an injury-specific manner. We hypothesized that diploid hepatocytes promote healing after acetaminophen (APAP)-induced liver injury.

**Approach & Results:** To study ploidy populations *in vivo*, we utilized mice with a lifelong liver-specific knockout of *E2f7*/*E2f8* (LKO), which are enriched in diploid hepatocytes (>70%) but otherwise normal. Control and LKO mice were treated with APAP (300 or 600 mg/kg), and injury was assessed over 0-96 hours. Although both groups sustained injury, LKO mice showed improved survival, lower serum liver enzyme levels, and reduced necrosis and DNA fragmentation, indicating resistance to APAP-induced injury. To determine if resistance was due to *E2f7/E2f8* loss or increased diploidy, we deleted *E2f7/E2f8* in adult hepatocytes (HKO), a model that does not alter ploidy. Injury was similar between controls and HKO, ruling out gene deletion as the protective factor. Transcriptomic and protein analyses revealed minimal baseline differences; however, following APAP treatment, LKO livers exhibited reduced JNK activation and less mitochondrial injury. Finally, APAP-treated wild-type hepatocytes exhibited a shift toward lower ploidy, supporting the idea that diploid cells are more resistant to injury.

**Conclusions:** Diploid hepatocytes resist APAP-induced liver injury through reduced JNK activation and mitochondrial damage. These findings highlight hepatocyte ploidy as a key determinant of injury response and suggest a protective role for diploid hepatocytes in promoting liver resilience and regeneration.

## Introduction

Most mammalian somatic cells are diploid, containing two copies of each chromosome set. However, certain cell types, including hepatocytes, undergo polyploidization, resulting in cells with multiple sets of chromosomes. The liver is a uniquely polyploid organ with polyploid hepatocytes comprising about 90% of the liver in adult mice and 20-50% in humans.^1,2^ Hepatic polyploidization primarily occurs during postnatal development through acytokinetic mitosis, where proliferating diploid hepatocytes undergo mitosis without cytokinesis, generating binucleate tetraploid daughter cells with diploid nuclei.^3–6^ These binucleate tetraploids can then enter subsequent cell cycles, completing mitosis with or without cytokinesis to produce a range of polyploid hepatocytes, including mononucleate and binucleate cells with higher DNA content.^7,8^ Hepatic polyploidy can also be reversed (ploidy reversal), generating lower-ploidy daughters that can then re-polyploidize.^1,9^ The regulation of hepatocyte polyploidization, ploidy reversal, and re-polyploidization, known as the ploidy conveyor, is controlled by a complex network of signaling pathways that influence cell cycle progression and cytokinesis.^3,4,10^

Key regulators of this process include the Epithelial-Splicing-Regulatory-Protein-2 (ESRP2)/miR-122 axis. ESRP2 enhances the biogenesis of miR-122, the most abundant microRNA in the liver.^11^ miR-122 expression spikes during postnatal development and promotes polyploidization by negatively regulating procytokinesis genes, including *RhoA*, *Mapre1*, and *Iqgap1*.^12^ Mice lacking miR-122 exhibit a significant reduction in polyploid hepatocytes.^13^ In contrast, excessive ploidy is restricted by the PIDDosome, a multiprotein complex activated by supernumerary centrosomes in polyploids. By inducing p21, the PIDDosome restricts hepatocyte proliferation and prevents hyperpolyploidization.^10,14–16^ PIDDosome loss leads to unrestrained polyploidy and increased tumor resistance.^14^ Among the most critical regulators of hepatic polyploidization are the atypical E2F family transcription factors, E2F7 and E2F8. These factors antagonize E2F1-mediated transcription and repress genes required for successful cytokinesis. Loss of E2F7/E2F8 in the postnatal liver shifts hepatocyte proliferation toward complete cytokinesis, blocking polyploidization.^17,18^ Liver-specific knockout of *E2f7/E2f8* (LKO) promotes the completion of cytokinesis during postnatal development, resulting in mature livers with hepatocytes that are predominately diploid.^17–20^ Despite this alteration in chromosomal content, *E2f7/E2f8* LKO mice maintain normal liver function into adulthood, making them a valuable model to investigate ploidy.

Although the mechanisms underlying hepatocyte polyploidization are well defined, the functions of distinct ploidy subpopulations remain unclear. Emerging data suggest that diploid and polyploid hepatocytes play distinct roles in liver physiology and disease. For example, excessive polyploidy is observed in metabolic dysfunction-associated steatotic liver disease (MASLD) and steatohepatitis (MASH), where hepatocytes show abnormal increases in mononucleated, high-ploidy cells.^21^ This pathological polyploidy, driven by oxidative stress and DNA damage, leads to mitotic inhibition and endoreplication.^21,22^ The consequences of such hyperpolyploidy are unclear, with potential roles in both promoting and protecting against disease. Polyploid hepatocytes can also undergo ploidy reversal, generating diploid or aneuploid progeny that enhance genetic diversity and promote adaptation to chronic liver injury. In mouse models of tyrosinemia, such aneuploid descendants clonally expand to repopulate the liver, conferring a selective advantage during injury.^1,23–25^ This highlights a role for polyploid-derived aneuploidy in providing adaptive genetic diversity. In hepatocellular carcinoma (HCC), ploidy plays a nuanced role. Most human and rodent HCCs are diploid, suggesting increased susceptibility of diploid hepatocytes to malignant transformation. Supporting this, polyploid hepatocytes can suppress tumor initiation, particularly when tumor suppressor genes are lost, by harboring extra copies of chromosomes that provide redundant tumor suppressor function.^26^ Once transformed, diploid hepatocytes tend to proliferate extensively, driving tumor growth.^19^ However, polyploid hepatocytes can undergo ploidy reduction, giving rise to diploid progeny that contribute to tumorigenesis.^9^ Moreover, polyploid HCCs have been identified in humans, particularly in tumors harboring *TP53* mutations.^2^

Our group recently demonstrated that hepatocyte ploidy influences proliferative capacity, with diploid hepatocytes exhibiting a distinct proliferative advantage over polyploids. In competitive repopulation assays, hepatocytes from highly diploid *E2f7/E2f8* LKO mice were co-transplanted with mostly polyploid wild-type (WT) hepatocytes into *Fah*^⁻/⁻^ recipients. Diploid-enriched hepatocytes consistently outcompeted polyploids during liver repopulation, indicating enhanced ability to proliferate under selective pressure. Consistent with these findings, we found that after partial hepatectomy in WT mice, diploid hepatocytes entered and progressed through the cell cycle more rapidly than polyploids. This enhanced proliferation by diploids was also observed in humans using retrospective radiocarbon birth dating, which revealed that diploid hepatocytes have a higher turnover rate than polyploids, suggesting that human hepatocyte proliferation is driven by diploids.^27^

We hypothesized that diploid-enriched livers would exhibit enhanced regeneration and protection after acute liver injury. To test this hypothesis, we used a well-characterized model of acetaminophen (APAP)-induced liver injury. APAP overdose is the leading cause of acute liver failure in the United States, accounting for nearly 50% of cases annually.^28^ Although APAP is safe at therapeutic doses, excessive intake leads to accumulation of its toxic metabolite, N-acetyl-p-benzoquinone imine (NAPQI).^29^ Under normal conditions, NAPQI is detoxified by conjugation with glutathione (GSH). However, during overdose, GSH stores are rapidly depleted, allowing NAPQI to form adducts with mitochondrial proteins, initiating oxidative stress and mitochondrial dysfunction.^30–32^ This triggers a cascade of cellular events, including c-Jun N-terminal kinase (JNK) activation, amplifying mitochondrial damage and necrotic cell death.^33,34^ While mild overdoses may be resolved through spontaneous liver regeneration, more severe injury can result in irreversible damage and progression to liver failure, where liver transplantation is the only treatment option.

Here, we investigated how hepatocyte ploidy affects susceptibility to APAP-induced liver injury by comparing diploid-enriched LKO and polyploid-enriched control mice. We found that diploid-enriched LKO were resistant to APAP-induced liver injury and initiated hepatocyte proliferation earlier than controls. This protective phenotype was associated with reduced JNK activation and diminished mitochondrial damage. Further, in WT hepatocytes, diploids were more resistant to APAP toxicity than polyploids. Together, these findings identify diploid hepatocytes as a protective subpopulation that promotes survival through stress signaling and regenerative dynamics.

## Materials and Methods

### Animals, Treatments, and Tissue Collection

All animal procedures were approved by the University of Pittsburgh Institutional Animal Care and Use Committee and followed NIH guidelines. WT C57BL/6J mice (Jackson Laboratory, #0664) and *E2f7/E2f8* floxed mice (from Drs. Alain de Bruin and Gustavo Leone)^17,18^ were used. Liver-specific *E2f7/E2f8* knockout (LKO) mice were of a mixed genetic background, predominantly FVB with floxed alleles for *E2f7/E2f8^loxP/loxP^*, R26R^mTmG/mTmG^, and Alb-Cre^Tg/0^ (Jackson Laboratory, #7676 and #3574).^35,36^ Control littermates lacked Cre (Alb-*Cre*^0^*^/^*^0^). Hepatocyte-specific knockouts (HKO) were generated by intraperitoneal (IP) injection in 2-month-old floxed mice with 1.25×10¹¹ AAV8-TBG-Cre viral particles; controls received AAV8-TBG-Null (gifted from James M. Wilson; Addgene, 107787-AAV8 and 105536-AAV8).

APAP (16 µg/µL in 0.9% saline) was administered IP at 300 or 600 mg/kg to 2.5-month-old overnight-fasted (∼16h) mice. Livers and blood were collected 0-96 hours post-injection. Liver tissue was snap-frozen; serum was isolated by centrifugation (10,000 rpm, 10 min, 4°C) and analyzed for alanine aminotransferase (ALT) and aspartate aminotransaminase (AST) by the University of Pittsburgh Medical Clinical Laboratory.

### Hepatocyte Isolation

Primary hepatocytes were isolated by a two-step collagenase perfusion, as previously published.^19^

### Ploidy analysis

Hepatocyte ploidy was evaluated by flow cytometry as previously described,^25^ using an LSR II flow cytometer (BD Biosciences) operated with BD FACSDiva™ Software v9.0. Data were analyzed and FACS plots generated using FlowJo v10.8.2 (FlowJo LLC).

### Protein Isolation and Western blotting

Lysates were prepared in T-PER buffer (ThermoFisher) with Halt protease/phosphatase inhibitors (ThermoFisher) and homogenized. Protein was quantified (Pierce PCA Assay; ThermoFisher), separated by SDS-PAGE (4-12% Bis-Tris gels), transferred to PVDF membranes (Millipore), blocked (Pierce Protein-Free buffer), and probed with primary and secondary antibodies. Detection used SuperSignal Pico substrate (ThermoFisher) and ChemiDoc Touch (Bio-Rad).

### Statistical analysis

Data are mean ± SEM. GraphPad Prism 10 was used. Two-group comparisons used unpaired two-tailed t-tests. Outliers were identified using Grubbs’ test (α = 0.01). Significance was set at P < 0.05.

For additional methods, please refer to the Supplemental Information.

## RESULTS

### Liver-Specific *E2f7/E2f8* Deletion Confers Resistance to APAP-Induced Hepatotoxicity

To investigate how hepatocyte ploidy influences acute liver injury, we used lifelong, Alb-Cre-driven liver-specific *E2f7/E2f8* knockout (LKO) mice. In contrast to control livers that are predominantly polyploid, with only 2-5% diploid hepatocytes, LKO livers have >70% diploid hepatocytes (Figure 1A,B, Supplemental Figure S1). To induce liver injury, control and LKO mice were fasted, administered APAP, and harvested 0-96 hours after overdose (Figure 1A). We first examined the response to 300 mg/kg APAP, previously characterized as a “regenerating dose” that induces injury followed by robust regeneration and spontaneous recovery.^37^ Following 300 mg/kg APAP, 25% of control mice died, while LKO mice survived completely (Figure 1C), suggesting resistance to APAP-induced damage. Biochemical analysis supported this observation, as serum levels of the liver injury biomarkers ALT and AST were significantly lower in LKO mice compared to controls, between 6 and 24 hours after injury (Figure 1D). Histological examination using H&E staining revealed widespread centrilobular necrosis in control livers, while LKO livers displayed markedly reduced necrosis from 6 to 72 hours (Figure 1E). Additionally, TUNEL staining showed reduced DNA fragmentation in LKO livers, further demonstrating APAP resistance by the LKO model (Figure 1F).

**Figure 1.**
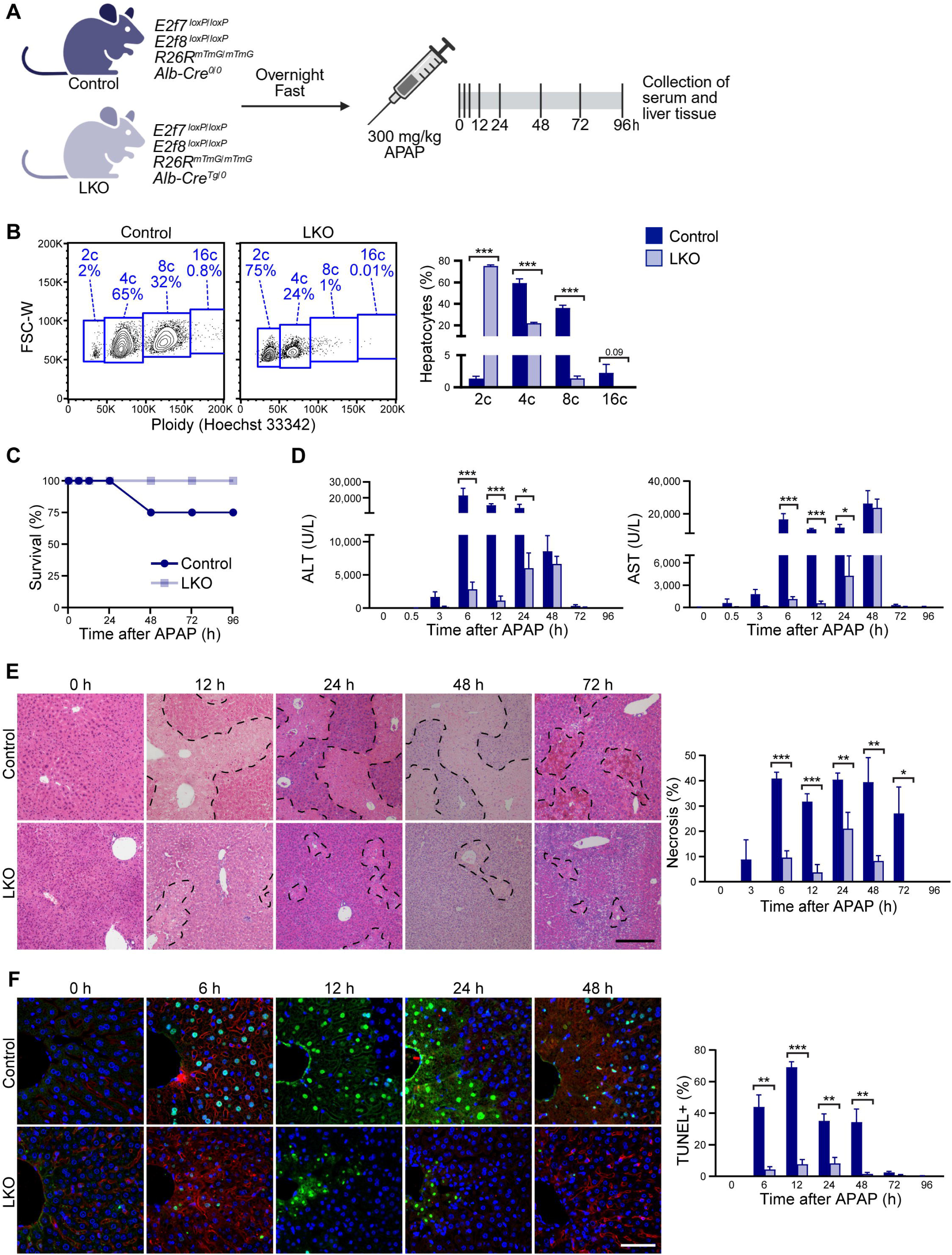
Liver-specific *E2f7/E2f8* deletion produces “polyploid knockout” mice resistant to APAP-induced hepatotoxicity. **(A)** Experimental scheme (Created with BioRender.com). **(B)** Freshly isolated hepatocytes from 3-month-old control and LKO mice were stained with Hoechst 33342 to determine cellular ploidy of viable hepatocytes by flow cytometry. Representative flow plots show the distribution of diploid (2c), tetraploid (4c), octaploid (8c), and hexadecaploid (16c) populations (n = 3-4/genotype; males and females). The full FACS gating strategy is shown in Supplemental Figure S1. **(C)** Survival curves of control and LKO mice following APAP overdose (n = 4-10/genotype/timepoint). **(D)** Levels of liver biomarkers ALT and AST in the serum (n = 4-9/genotype/timepoint). **(E)** Quantification of necrosis by H&E staining (n = 4-9/genotype/timepoint). Scale bar = 200 µM. **(F)** TUNEL staining showing DNA fragmentation (green) and nuclear staining with Hoechst 33342 (blue) (n = 4-9/genotype/timepoint). Scale bar = 50 µM. Representative images, plots, and quantification results are shown. Graphs show mean ± SEM. *P < 0.05; **P < 0.01; ***P < 0.001.

To test whether LKO mice retained this protective phenotype under more stringent conditions, we next challenged them with 600 mg/kg APAP, a “non-regenerating dose” previously shown to cause sustained hepatocellular injury, impaired regenerative responses, and poor survival outcomes.^37^ At this higher dose, the mortality of control mice increased dramatically, with 91% dying within 72 hours (Supplemental Figure S2A). Strikingly, 53% of LKO mice survived the overdose, highlighting a significant survival advantage even in a nonregenerative context. Among the surviving animals, ALT and AST levels were comparable between genotypes, while necrosis and DNA fragmentation were reduced in LKO mice at 48 and 24 hours, respectively (Supplemental Figure S2B-D). The injury observed in the control cohort may be underestimated, as only 9% of mice survived to 72 hours, and samples were collected from these rare survivors. Together, these results demonstrate that diploid-enriched LKO mice significantly resist APAP-induced liver injury across both regenerative and nonregenerative models.

### Diploid-rich LKO Livers Exhibit Enhanced Proliferation

Considering our earlier work that diploid hepatocytes are primed for proliferation and regeneration, we next examined whether LKO mice exhibit altered regenerative responses following 300 mg/kg APAP injury.^19^ Whole liver lysates were analyzed for key proteins involved in proliferative signaling. Notably, levels of active β-CATENIN (non-phosphorylated at Ser45), which directly promotes expression of the G1/S cell cycle regulator Cyclin D1, were comparable between groups at baseline and up to 24 hours post-injury. However, by 48 hours, LKO mice showed significantly elevated active β-CATENIN compared to controls, suggesting that diploid hepatocytes in LKO livers initiate cell cycle entry earlier (Figure 2A, pooled samples; Supplemental Figure S3A, individual replicates). This trend is mirrored by Cyclin D1 and PCNA, which increase by 48 hours after APAP overdose, indicating compensatory regeneration occurs, with significantly increased expression in the LKO at 48 hours. By 96 hours, the pattern is reversed, with control mice displaying higher Cyclin D1 and PCNA, indicating a delayed but robust regenerative response (Figure 2B,C, Supplemental Figure S3B). To assess whether differences in β-CATENIN activation were due to altered degradation, we also examined phosphorylated β-CATENIN (Ser33/37/Thr41), which targets the protein for proteasomal degradation, and found no differences between groups across timepoints.

**Figure 2.**
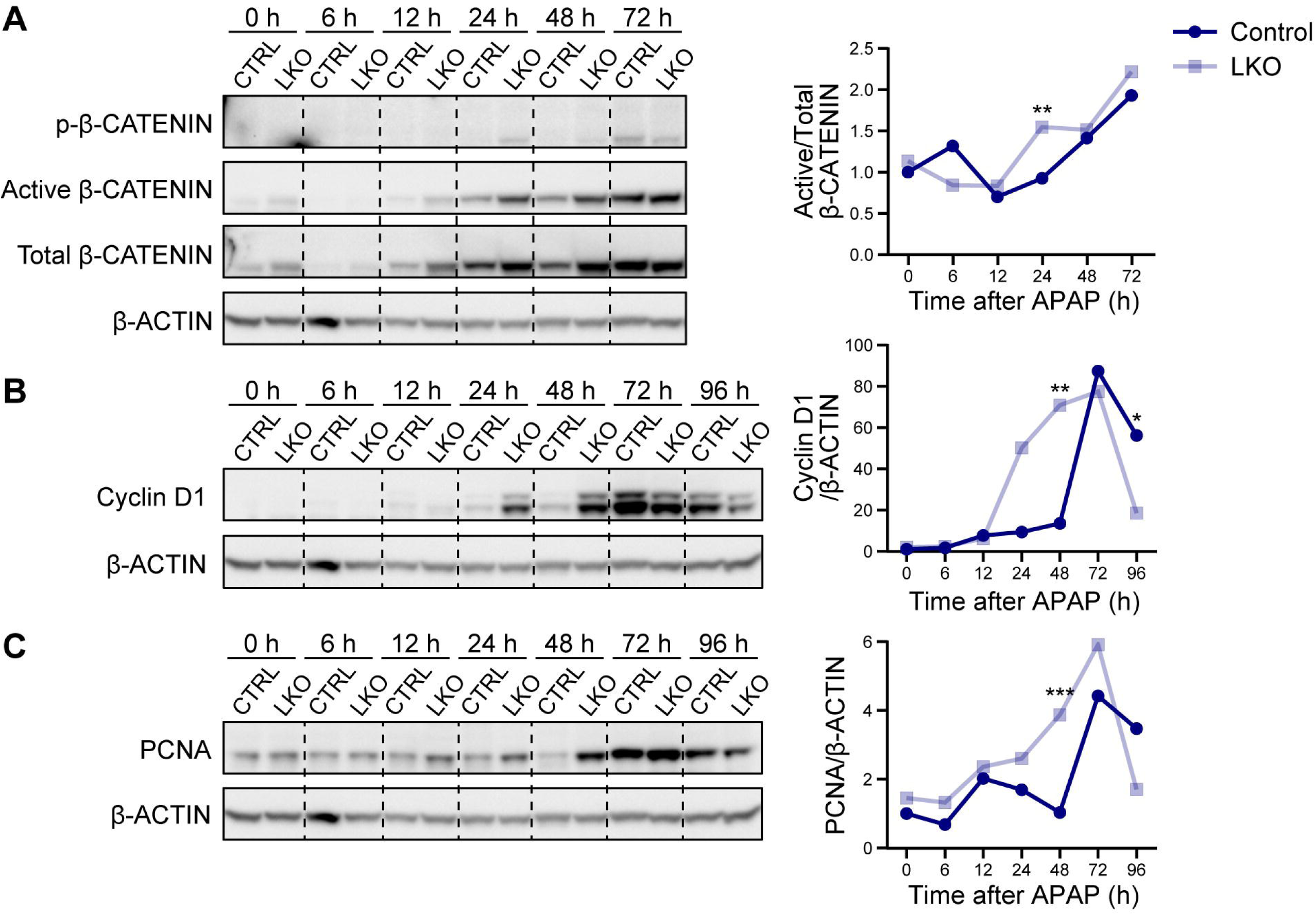
LKO hepatocytes display enhanced proliferation APAP overdose. **(A-C)** Western blotting of whole liver lysates collected from control and LKO mice 0-96 hours after 300 mg/kg APAP overdose, showing expression of proteins involved in proliferation: **(A)** active β-CATENIN relative to total β-CATENIN (pooled samples; n = 3/genotype/timepoint), **(B)** Cyclin D1 relative to β-ACTIN (pooled samples; n = 3/genotype/timepoint), **(C)** PCNA relative to β-ACTIN (pooled samples; n = 3/genotype/timepoint). Shown are representative blots and quantification results normalized to the 0 hour control, which is set to 1. The significance values are based on western blot analysis of individual samples shown in Supplemental Figure S2 (n = 3-5/genotype/timepoint). *P < 0.05; **P < 0.01; ***P < 0.001.

Despite sustaining markedly less liver injury after APAP injection, LKO mice expressed proliferation markers at levels comparable to or greater than those of controls. This finding suggests that diploid hepatocytes in LKO livers not only resist APAP-induced injury but also initiate regeneration more rapidly, consistent with our previous findings that LKO livers and diploid hepatocytes enter and progress through the cell cycle faster than WT hepatocytes.^19^

### *E2f7/E2f8* Deletion Alone Does Not Confer Resistance to APAP-Induced Injury

To determine whether APAP resistance observed in LKO mice was due to gene expression changes associated with *E2f7*/*E2f8* deletion or the enrichment of diploid hepatocytes in the LKO model, we generated an inducible hepatocyte-specific knockout (HKO) model by injecting adult mice (containing loxP sites in *E2f7* and *E2f8*) with AAV8-TBG-Cre to delete *E2f7/E2f8*. Control animals received AAV8-TBG-Null control virus. Because adult livers are already predominantly polyploid, this approach enables the deletion of *E2f7/E2f8* without altering hepatocyte ploidy. Two weeks after AAV injection, mice were fasted, administered 300 mg/kg APAP, and harvested at multiple timepoints (Figure 3A).

**Figure 3.**
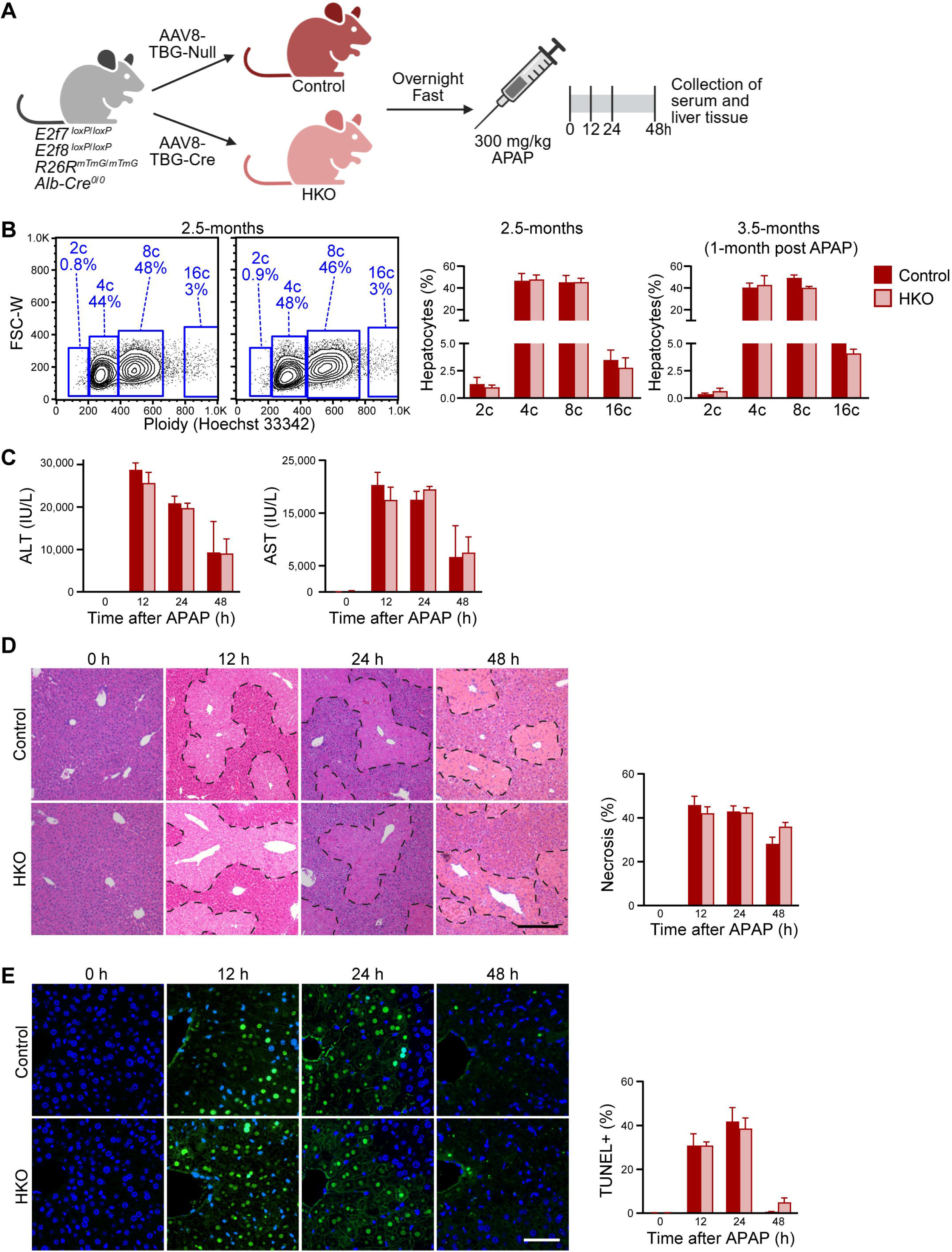
Loss of *E2f7/E2f8* in adults with normal ploidy does not protect against APAP-induced damage. **(A)** Experimental scheme (Created with BioRender.com). **(B)** Ploidy analysis by flow cytometry of freshly isolated hepatocytes from 2.5-month-old control and HKO mice at baseline, and from 3.5-month-old control and HKO mice one month after 300 mg/kg APAP treatment and recovery. (n = 3/genotype/timepoint; males). **(C)** Levels of liver biomarkers ALT and AST in the serum (n = 3-6/genotype/timepoint). **(D)** Quantification of necrosis by H&E staining (n = 3-6/genotype/timepoint). Scale bar = 200 µM. **(E)** TUNEL staining showing DNA fragmentation (green) and nuclear staining with Hoechst 33342 (blue) (n = 3-6/genotype/timepoint). Representative images, plots, and quantification results are shown. Graphs show mean ± SEM. *P < 0.05; **P < 0.01; ***P < 0.001.

To evaluate the HKO model, we first examined the R26R-mTmG reporter by fluorescent imaging. Two weeks after AAV8 injection, all hepatocytes in control livers were tdTomato^+^, while >99% of hepatocytes in HKO livers were GFP^+^, indicating efficient Cre-mediated recombination (Supplemental Figure S4A). Next, to assess baseline transcriptomic differences between control and HKO mice following AAV8-TBG-Cre injection, we performed RNA sequencing on whole livers at 0 hours. Knockout of *E2f8* was confirmed by evaluating gene counts for functional exons 3 and 4. *E2f8* expression was significantly reduced in HKO mice two weeks after AAV8-TBG-Cre injection, indicating efficient Cre-mediated deletion (Supplemental Figure S4B). *E2f7* is lowly, if at all, expressed in adult livers and was below the threshold for detection. Transcriptomic analysis revealed 320 differentially expressed genes between HKO and control, suggesting relatively modest changes at baseline (Supplemental Figure S4C; Supplemental Table S1). Finally, ploidy analysis demonstrated that both control and HKO livers remained predominantly diploid at baseline (Figure 3B). Moreover, one month following APAP treatment, when livers had fully recovered from APAP overdose, the HKO ploidy spectrum remained equivalent to controls, indicating that loss of *E2f7*/*E2f8* in adults does not affect ploidy even after extensive compensatory liver regeneration (Figure 3B).

Biochemical and histological analyses revealed no significant differences in liver injury between control and HKO mice after overdose. Serum ALT and AST levels were comparable at all time points (Figure 3C), and H&E and TUNEL staining showed similar levels of necrosis and DNA fragmentation, respectively (Figure 3D,E). Additionally, key proliferative signaling markers, active β-CATENIN, Cyclin D1, and PCNA, were expressed similarly between HKO and controls at most time points, with only minor differences observed (Supplemental Figure S4D,E). In contrast to the LKO model, HKO mice showed no alterations in ploidy, injury, or regenerative signaling after APAP. These findings indicate that the resistance to APAP-induced toxicity observed in LKO mice is independent of *E2f7/E2f8* deletion and is likely attributed to the high percentage of diploid hepatocytes.

### APAP Metabolism Machinery Is Conserved in the LKO Model

To identify the mechanism driving APAP resistance by diploid hepatocytes, we turned to the LKO model and assessed whether differences in APAP metabolism could account for the phenotype. Bulk RNA-seq of livers from control and LKO mice that were fasted but untreated (0 hour timepoint) revealed few differentially expressed genes, consistent with previously published findings on this model (Supplemental Figure S5A, Supplemental Table S2).^20^ Given the minimal transcriptional differences observed, we next examined protein levels of enzymes involved in APAP metabolism to determine whether post-transcriptional regulation could account for the observed phenotype. Protein expression of cytochrome P450 enzymes CYP2E1 and CYP1A2, key mediators of APAP bioactivation to the hepatotoxic metabolite NAPQI, was unchanged between LKO and control livers at baseline (Figure 4A). We next examined hepatic GSH levels, the major detoxifying molecule that neutralizes the toxic APAP metabolite NAPQI. GSH concentrations at 0 hour and 30 minutes after 300 mg/kg APAP were similar between groups, indicating that LKO and control mice have comparable capacity for detoxification (Figure 4B). Additionally, APAP-protein adduct formation was similar between LKO and controls at 0.5, 6, and 12 hours after 300 mg/kg APAP (Figure 4C). Thus, APAP metabolism and detoxification are intact in both groups.

**Figure 4.**
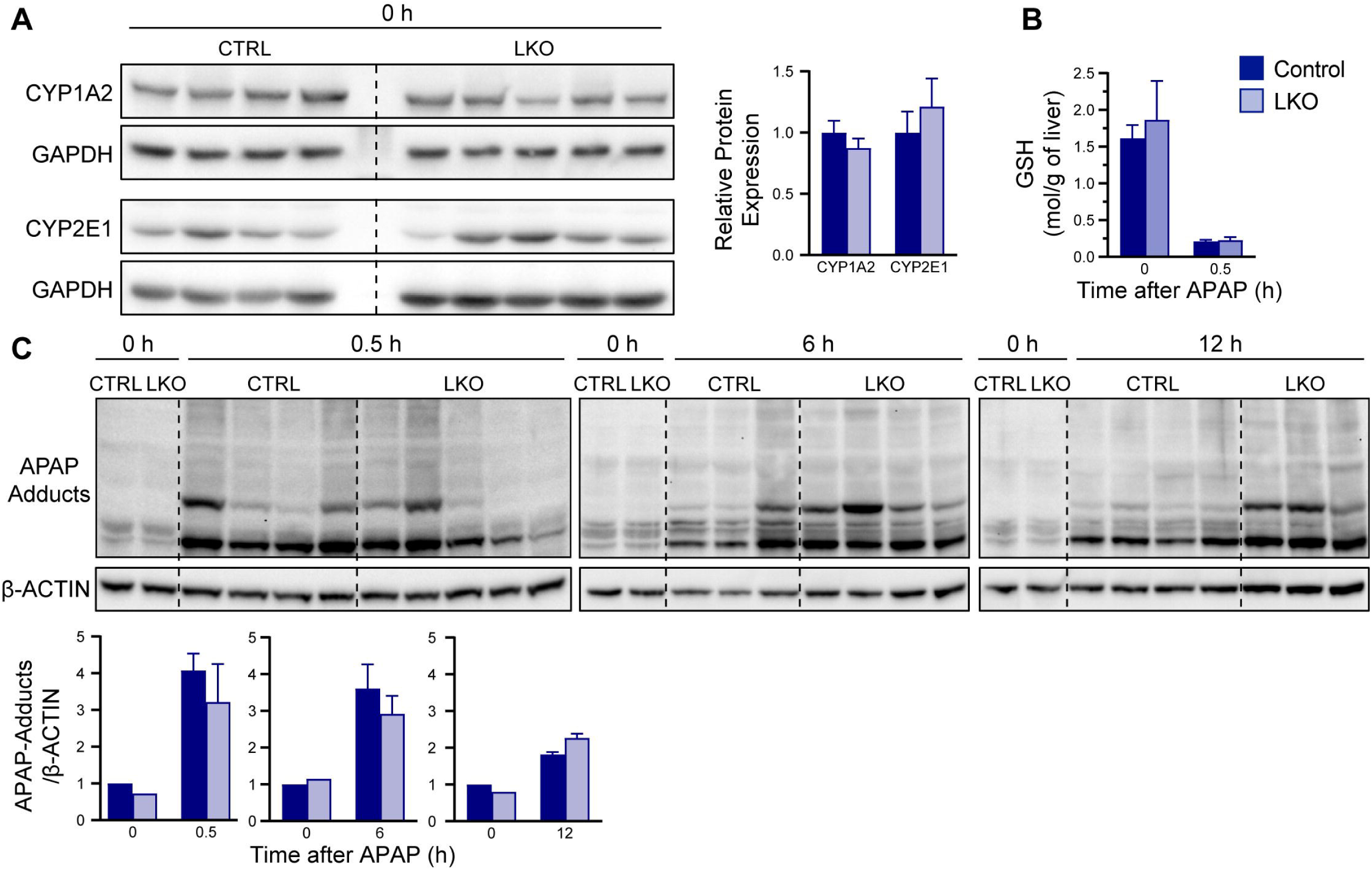
APAP metabolism machinery is conserved in the LKO model. **(A)** Expression levels of key APAP-metabolizing enzymes CYP2E1 and CYP1A2 in control and LKO mice at baseline (n = 4-5/genotype/timepoint). **(B)** Baseline hepatic GSH levels and its depletion 0.5 hours after 300 mg/kg APAP overdose in control and LKO mice (n = 4-6/genotype/timepoint). (C) Formation of APAP-protein adducts at 0, 0.5, 6, and 12 hours following 300 mg/kg APAP (n = 3-5/genotype/timepoint). Graphs show mean ± SEM. *P < 0.05; **P < 0.01; ***P < 0.001.

Although few gene expression differences were observed at baseline, bulk RNA-seq analysis at 12, 24, 48, 72, and 96 hours after APAP overdose revealed divergent transcriptional responses between control and LKO mice. Principal component analysis showed that samples from both groups are clustered tightly together at 0 hours, consistent with minimal transcriptional differences under baseline conditions (Supplemental Figure S5B). However, following APAP injury, the two groups separated along principal components 1 and 2, indicating distinct transcriptional trajectories. Control livers followed a broad arc, with gene expression divergence that peaked between 12 and 24 hours. In contrast, LKO livers show a more limited change in gene activity that remained distinct from controls for most of the time course. By 96 hours, the two groups moved closer together toward the 0 hour time point again, suggesting resolution of injury.

Broad, temporal changes in gene expression were observed between control and LKO mice after APAP overdose (Supplemental Figure S5C, Supplemental Table S3). For example, genes in cluster 1, which were rapidly upregulated in control livers within 12 hours of APAP administration, showed delayed or absent induction in LKO livers. In contrast, genes in cluster 3 that were induced later (72-96 hours) in controls appeared to be activated earlier in LKO livers, suggesting a dysregulated injury response. Strikingly, key differentially expressed genes included those involved in cellular injury, oxidative stress, mitochondrial function, and proliferation (Supplemental Figure S5D, Supplemental Table S4). One key example is *Cdkn1a,* which encodes p21, a regulator of cell cycle arrest and DNA damage response. In controls, *Cdkn1a* was strongly upregulated and remained elevated for a longer duration than in LKO, consistent with a more sustained injury response. Ingenuity Pathway Analysis (IPA) identified the activation of WNT/β-catenin signaling in LKO livers (Supplemental Figure S5E), aligning with increased protein levels of active β-CATENIN and its downstream target Cyclin D1 (Figure 2, Supplemental Figure S3). Sirtuin signaling, a pathway involved in oxidative stress responses and previously linked to APAP toxicity, was also predicted to be activated in LKO mice.^38–42^ However, protein levels of SIRT1, SIRT3, and SIRT6 at 0 and 6 hours post-APAP were similar between groups (Supplemental Figure S5F), suggesting that regulation of Sirtuins does not account for the observed hepatoprotection.

Together, these results demonstrate that LKO mice have intact APAP metabolism and detoxification capacity, indicating that their resistance to APAP-induced liver injury is not due to differences in early drug metabolism. Instead, the findings point to altered injury and recovery responses in LKO livers. This suggests that diploid and polyploid hepatocytes respond differently to stress, with diploids potentially mounting a more protective or regenerative transcriptional response following APAP injury.

### Blunted JNK Activation Mediates Hepatoprotection in LKO Mice

Considering the pattern of attenuated expression of genes related to cell death and injury combined with preserved mitochondrial function in the LKO mice, we next examined the JNK pathway, a key driver of APAP-induced hepatotoxicity. JNK activation, especially mitochondrial translocation of its phosphorylated form (pJNK), exacerbates mitochondrial oxidative stress and promotes necrotic cell death.^33,43,44^ Western blot analysis revealed significantly lower levels of pJNK in LKO livers compared to controls at multiple timepoints following APAP overdose (Figure 5A, pooled samples; Supplemental Figure S6, individual replicates). Consistent with this finding, mitochondrial fractions isolated from livers 6 hours post-APAP showed reduced accumulation of total and phosphorylated JNK in LKO mice relative to controls (Figure 5B). To assess downstream effects on mitochondrial integrity, we measured cytosolic levels of apoptosis-inducing factor (AIF), a mitochondrial endonuclease released during stress-induced necrosis.^44–46^ Six hours post-APAP, LKO livers had markedly lower levels of cytoplasmic AIF than controls (Figure 5C), suggesting reduced mitochondrial dysfunction. This protection aligns with transcriptional data showing that genes and pathways associated with oxidative stress and mitochondrial health are differentially regulated in LKO livers (Supplemental Figure S5D,E). Together, these data indicate that diploid-enriched LKO livers suppress JNK activation and mitochondrial translocation, which helps maintain mitochondrial integrity and reduces necrotic cell death after APAP-induced injury.

**Figure 5.**
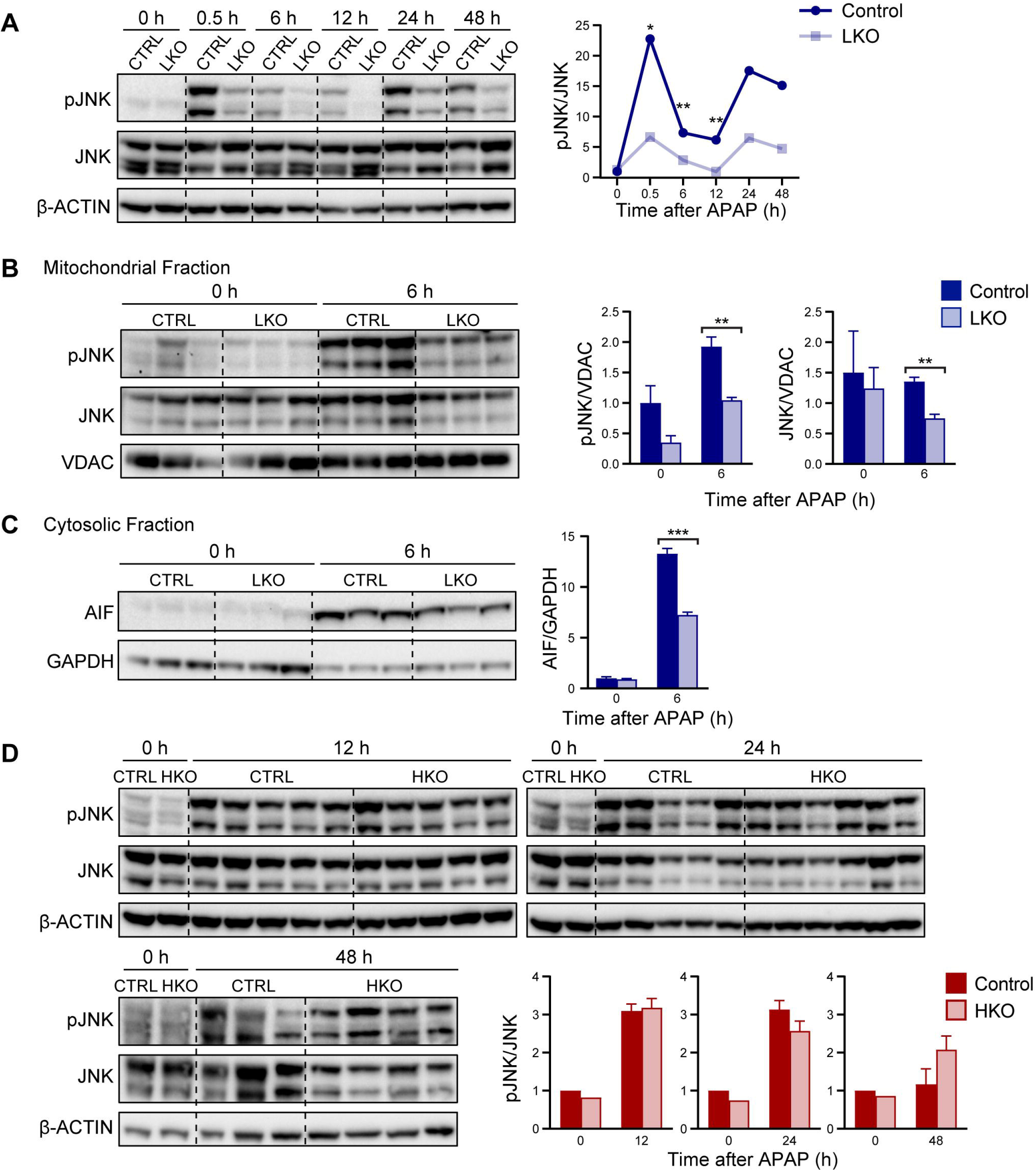
APAP resistance in the LKO model is mediated by JNK activation. **(A)** Expression of pJNK relative to total JNK in whole liver lysates from control and LKO mice 0-48 hours after 300 mg/kg APAP overdose. Samples are pooled (n=3/genotype/timepoint). The significance values are based on western blot analysis of individual samples shown in Supplemental Figure S6. **(B)** Expression of pJNK and total JNK relative to VDAC in the mitochondrial fraction of control and LKO livers 0 and 6 hours after APAP (n = 3/genotype/timepoint). **(C)** Expression of cytosolic AIF in control and LKO livers 0 and 6 hours after APAP (n = 3-5/genotype/timepoint). (**D**) Expression of pJNK relative to total JNK in HKO and control livers 6 hours after APAP (n = 3-4/genotype). Shown are representative blots and quantification results normalized to the 0 hour control, which is set to 1. Graphs show mean ± SEM. *P < 0.05; **P < 0.01; ***P < 0.001.

To distinguish whether reduced JNK activation arises from increased diploidy or from deletion of *E2f7/E2f8*, we evaluated pJNK levels in HKO mice following APAP treatment. Unlike LKO mice, HKO showed no reduction in JNK activation compared to controls (Figure 5D), suggesting that reduced JNK activation in LKO livers is not a direct consequence of *E2f7*/*E2f8* loss but is instead likely driven by the enrichment of diploid hepatocytes. Taken together, these data suggest that reduced JNK activation and diminished mitochondrial stress signaling drive resistance to APAP toxicity observed in diploid hepatocytes of LKO mice.

### Highly polyploid hepatocytes are sensitive to APAP toxicity

To determine whether the APAP resistance observed in diploid-enriched LKO mice is a general feature of diploid hepatocytes, we evaluated APAP sensitivity in WT hepatocytes *in vitro*. Primary hepatocytes were isolated from adult C57BL/6J mice, treated with increasing concentrations of APAP (0, 5, and 10 mM), and assessed after 24 hours (Figure 6A). Microscopy revealed a clear dose-dependent increase in cellular injury, including hepatocyte rounding, increased lipid droplet accumulation, and increased cell death with rising APAP concentrations (Figure 6B). To quantitatively assess cell death, hepatocytes were stained with a fixable viability dye (FVD) and Hoechst 33342, allowing for the simultaneous measurement of viability and ploidy using flow cytometry. As expected, APAP treatment led to a dose-dependent increase in cell death, with the proportion of viable hepatocytes decreasing from 58% at 0 mM APAP to 17% at 10 mM (Figure 6C).

**Figure 6.**
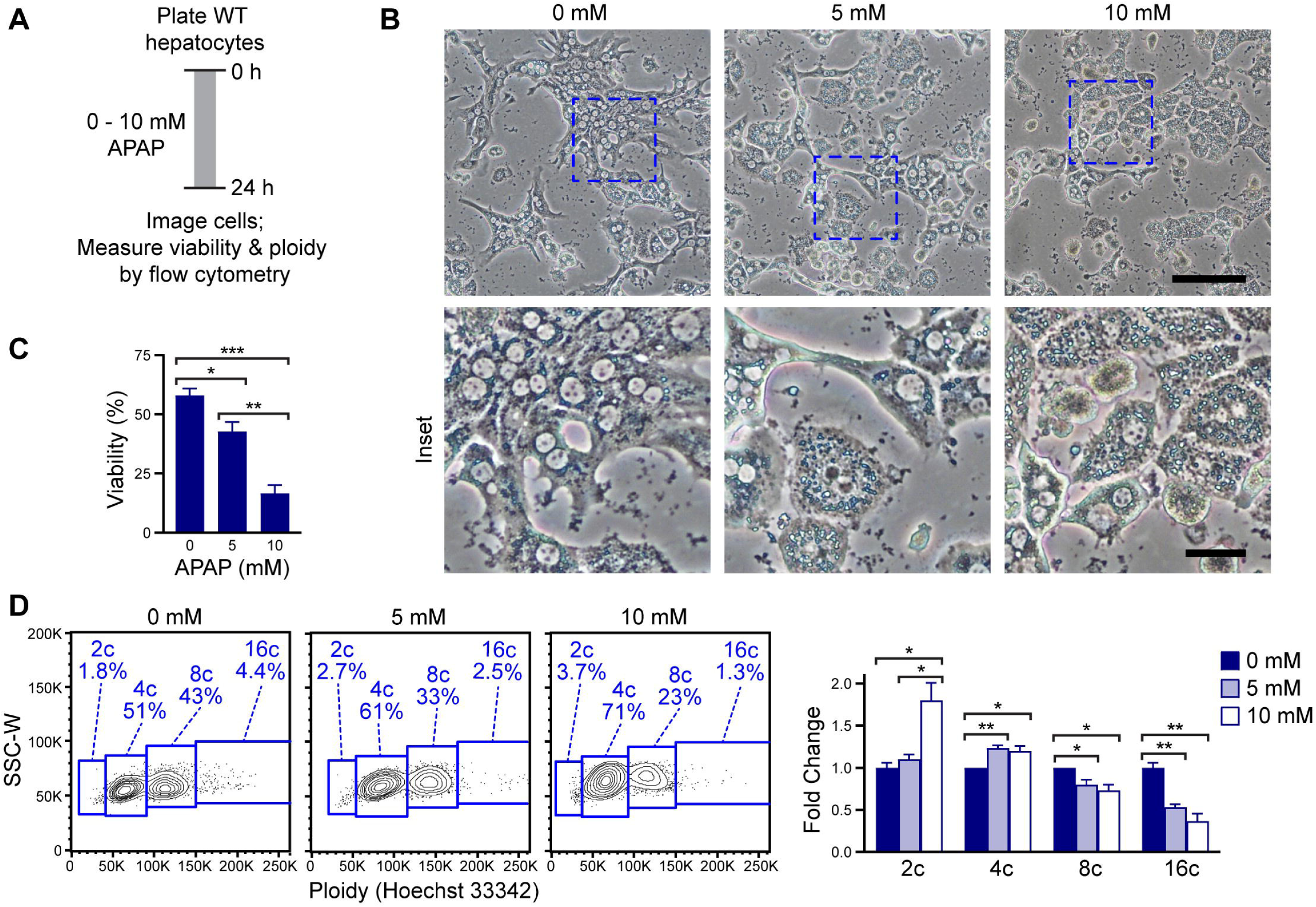
Highly polyploid hepatocytes are more susceptible to APAP-induced cell death. **(A)** Schematic of *in vitro* APAP toxicity assay. Primary hepatocytes were isolated from 2-3-month-old WT mice (male), treated with increasing doses of APAP (0, 5, and 10 mM) for 24 hours. **(B)** Representative images showing APAP-induced dose-dependent hepatocyte injury at 24 hours; higher-magnification insets highlight cellular detail (main image scale bar = 100Lµm; inset scale bar = 25Lµm). **(C)** Quantification of cell death by FVD staining. **(D)** Flow cytometry plots showing relative ploidy distribution (2c, 4c, 8c, 16c) within viable (FVD negative) hepatocytes. Data are shown as the fold change of each ploidy population, where 0 mM APAP was set to 1. Data shown have n = 3 internal replicates per condition and are representative of 4 independent experiments. Graphs show mean ± SEM. *P < 0.05; **P < 0.01; ***P < 0.001.

To investigate whether ploidy status influences susceptibility to APAP-induced cell death, we analyzed the distribution of hepatocyte ploidy in viable populations. In untreated cultures (0 mM), the expected distribution of 2c, 4c, 8c, and 16c hepatocytes was observed. However, following treatment with 10 mM APAP, when substantial cell death was present, there was a consistent decrease in the proportion of live 8c and 16c hepatocytes and an enrichment of 2c and 4c populations. For instance, treatment with 10 mM APAP led to a reduction in viable high-ploidy hepatocytes, with 8c cells decreasing from 40% to 29%, and 16c cells from 3.5% to 1.5%. In contrast, low-ploidy populations increased, with 2c hepatocytes doubling from 2% to 4% and 4c cells rising from 51% to 63% (Figure 6D). Together, these data demonstrate that hepatocyte sensitivity to APAP-induced toxicity is influenced by ploidy status, with low-ploidy cells less susceptible to injury and death. These *in vitro* findings support the *in vivo* observation that diploid-enriched livers are more resistant to APAP and suggest that reduced ploidy states may confer a protective advantage in the context of acute hepatotoxic stress.

## Discussion

The liver is characterized by extensive hepatocyte polyploidy, yet the functional significance of different ploidy states remains incompletely understood. Here, we demonstrate that hepatocyte ploidy modulates the liver’s response to acute drug-induced injury. Using a liver-specific *E2f7*/*E2f8* knockout model (LKO) enriched in diploid hepatocytes, along with WT hepatocytes, we show that diploid hepatocytes are unexpectedly resistant to APAP-induced toxicity and initiate more rapid compensatory regeneration than their polyploid counterparts.

Diploid-enriched LKO mice exhibited reduced serum liver enzyme levels, diminished histological damage, and lower DNA fragmentation following APAP overdose. Despite sustaining less injury, LKO livers mounted an earlier and more robust regenerative response, evidenced by activation of WNT/β-catenin signaling and increased expression of Cyclin D1 and PCNA. These findings align with prior reports^19^ that diploid hepatocytes are primed for proliferation and extend this concept by revealing a previously unrecognized role in injury resistance. To determine whether this protective phenotype resulted from diploid enrichment or from direct effects of *E2f7/E2f8* loss, we generated an inducible hepatocyte-specific knockout (HKO) model in which gene deletion occurs in adult, polyploid livers. HKO mice exhibited APAP-induced liver injury comparable to controls, indicating that the protection observed in LKO mice is not due to loss of *E2f7/E2f8* alone but rather to increased diploidy. To further validate the link between ploidy and APAP resistance, we conducted assays with WT hepatocytes and assessed viability across ploidy populations. Highly polyploid cells (8c, 16c) were significantly more sensitive to APAP-induced death than low-ploidy cells (2c, 4c), confirming that ploidy directly influences toxicity.

Mechanistically, resistance in LKO livers was not due to altered APAP metabolism. Expression of key cytochrome P450 enzymes (CYP2E1/CYP1A2), GSH depletion, and APAP-protein adduct formation were similar between LKO and controls. Instead, transcriptomic profiling revealed a distinct injury response in LKO livers, marked by reduced activation of injury-related genes and earlier induction of regeneration, including WNT/β-catenin signaling and Cyclin D1 expression. Strikingly, there were profound differences in JNK signaling between LKO and control livers (Figure 7A). JNK is a well-established driver of APAP-induced hepatocyte necrosis, and its suppression was a key distinguishing feature of the diploid-enriched response. In LKO, phosphorylated JNK levels and mitochondrial translocation were decreased, along with cytosolic release of AIF, a marker of mitochondrial injury. These findings align with previous reports showing that pharmacologic inhibition and germline knockout of JNK protects against APAP toxicity^34,44,47–49^, support a model in which diploid hepatocytes resist injury through suppression of JNK-mediated cell death.

**Figure 7.**
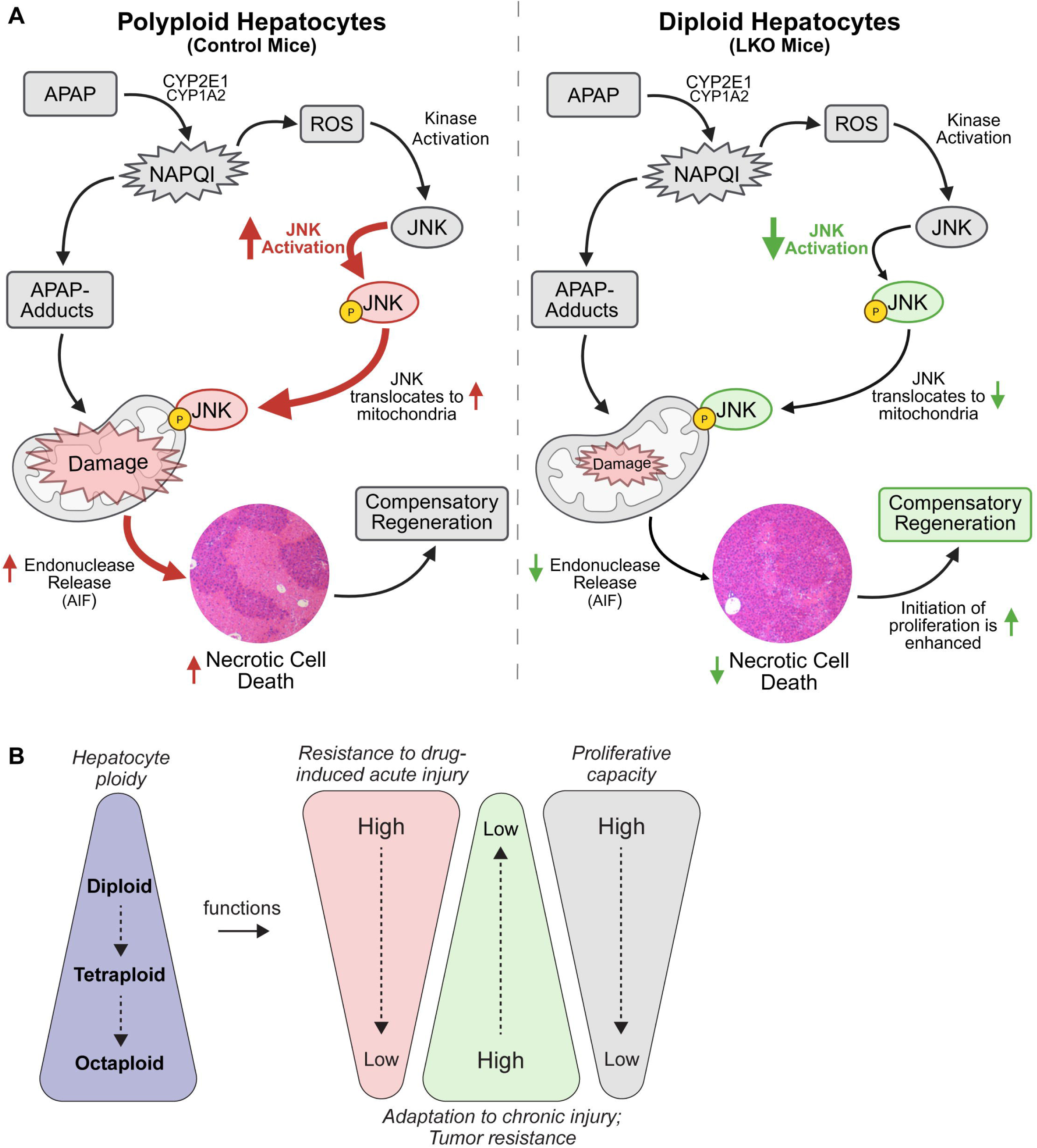
Summary of the ploidy-dependent response to APAP overdose in the liver. **(A)** Schematic summary comparing how polyploid (control) and diploid-enriched (LKO) livers respond to APAP overdose (Created with Biorender.com). Both cell types metabolize APAP through cytochrome P450 enzymes (CYP2E1, CYP1A2), generating the toxic intermediate NAPQI and forming APAP-protein adducts. In polyploid hepatocytes, this leads to oxidative stress, JNK activation and mitochondrial translocation, release of AIF, and necrotic cell death. In contrast, diploid hepatocytes suppress JNK activation and mitochondrial injury, limiting cell death. **(B)** Model illustrating context-dependent advantages of hepatocyte ploidy states. Diploid hepatocytes exhibit enhanced resistance to acute toxic injury, such as APAP overdose, and demonstrate a greater capacity for proliferation. In contrast, polyploid hepatocytes support tumor suppression and adaptation to chronic injury through mechanisms such as aneuploidy tolerance and metabolic buffering. This functional specialization enables the liver to respond flexibly across diverse injury contexts.

Spatial zonation within the liver lobule is a critical consideration in APAP-induced injury, which targets pericentral hepatocytes with high CYP2E1 expression. If polyploid hepatocytes were selectively localized to this region, their loss could explain greater injury in polyploid-rich livers. However, prior work shows that polyploid hepatocytes are not exclusively zonated and that LKO mice retain normal hepatic architecture. ^2,8,19,50,51^ Thus, the protective phenotype in LKO mice is unlikely to result from altered zonation.

Our results also align with findings in *Mir122* knockout mice, which show increased proportions of diploid hepatocytes (like LKO mice) and APAP resistance.^13,52^ While those effects were attributed to reduced CYP2E1/CYP1A2 expression, diploid enrichment may also contribute. The proliferative advantage of diploid hepatocytes may also support their resistance to APAP injury. Proliferation has been linked to induction of efflux transporter MRP4, which transports byproducts of cell injury, promoting resistance to liver injury after multiple rounds of APAP overdose.^53^ These findings suggest that diploid-enriched livers are better equipped for regeneration, detoxification, and stress resilience, supporting a model where increased diploidy enhances liver recovery during acute toxic stress.

Prior studies have suggested that polyploidy enhances cellular functional capacity, potentially through increased gene dosage^54^ and modest transcriptional changes. For example, Lu et al. identified small changes in gene expression between ploidy states – such as increased *Nr1i3* and *Ccne2*, and decreased *Gas2* – which may support enhanced stress response.^55^ Mietten et al. found that higher ploidy correlates with reduced mitochondrial gene expression, possibly conferring protection from oxidative stress.^56^ Despite these potential advantages, our findings reveal that diploid hepatocytes exhibit superior stress tolerance following APAP injury, mediated by suppressed JNK signaling and enhanced regeneration. Diploid hepatocytes may also better preserve mitochondrial function or redox balance under stress. Although baseline mitochondrial gene expression was similar between groups, post-injury transcriptional differences suggest diploid hepatocytes may maintain mitochondrial integrity more effectively.

Collectively, these findings support a model in which hepatocyte ploidy states confer distinct, context-dependent advantages. Polyploid hepatocytes may promote tumor suppression and adaptation to chronic liver injury through mechanisms like aneuploidy generation and metabolic buffering.^1,6,23^ In contrast, diploid hepatocytes appear better equipped for acute toxic injury, resisting necrosis and initiating regeneration. This division of labor may allow flexible responses to diverse injury contexts (Figure 7B). While human data remain limited, multiple studies report wide variability in hepatocyte ploidy among individuals, with estimates ranging from 20-50% polyploidy depending on methodology.^2,57–59^ This individual variability raises the possibility that ploidy composition could influence clinical outcomes. We speculate that higher diploid content may confer resilience to APAP toxicity and that hepatocyte ploidy profiling could help predict injury susceptibility.

In conclusion, this study establishes hepatocyte ploidy as a key determinant of susceptibility to APAP-induced liver injury. Diploid hepatocytes resist necrotic cell death through suppression of JNK signaling and initiate robust regeneration, enabling recovery under toxic stress. This work not only defines a mechanistic basis for diploid resilience but also supports a broader model in which ploidy states specify distinct functional roles relevant to human liver disease.

## Acknowledgements

We thank Gustavo Leone (Medical College of Wisconsin) and Alain deBruin (Utrecht University) for sharing liver-specific *E2f7/E2f8* knockout mice. Flow cytometry data were generated in the University of Pittsburgh Unified Flow Cytometry Core Facility (RRID:SCR_025102); special thanks to Nan Sheng and Dewayne Falkner for flow cytometry expertise.

## List of Abbreviations

AIF: apoptosis inducing factor
ALT: alanine aminotransferase
APAP: acetaminophen
AST: aspartate aminotransferase
FACS: fluorescence-activated cell sorting
FBS: fetal bovine serum
FVD: fixable viability dye
GFP: green fluorescent protein
GSH: glutathione
H&E: hematoxylin and eosin
HKO: hepatocyte-specific *E2f7E2f8* knockout
HCC: hepatocellular carcinoma
IP: intraperitoneal
IPA: Ingenuity pathway analysis
JNK: c-Jun N-terminal kinase
LKO: liver-specific *E2f7/E2f8* knockout
MASLD: metabolic dysfunction-associated steatotic liver disease
MASH: metabolic dysfunction-associated steatohepatitis
NAPQI: N-acetyl-p-benzoquinone imine
PCA: principal component analysis
PCNA: proliferating cell nuclear antigen
PIDDosome: p53-induced protein with a death domain (PIDD) signaling complex
SEM: standard error of mean
TUNEL: terminal deoxynucleotidyl transferase dUTP nick end labeling
WT: wild-type

