## Supplemental Information for "Diploid Hepatocytes Resist Acetaminophen-Induced Liver Injury Through Suppressed JNK Signaling"

**Supplemental Experimental Procedures**

**Supplemental Figures**

| **Supplemental Figure S1.** | Flow cytometry gating strategy for analysis of hepatocyte ploidy distribution. |
| --- | --- |
| **Supplemental Figure S2.** | LKO mice resist lethal dose of APAP (600 mg/kg). |
| **Supplemental Figure S3.** | LKO mice show enhanced proliferation at 48 hours. |
| **Supplemental Figure S4.** | Expression in the HKO model. |
| **Supplemental Figure S5.** | Gene and protein expression in LKO livers. |
| **Supplemental Figure S6.** | Reduced JNK activation in LKO mice after APAP overdose. |

**Supplemental Experimental Procedures**

**Tissue processing and staining:** Liver tissues were fixed in 10% neutral-buffered formalin, embedded in paraffin, and sectioned at a thickness of 4 μm. Hematoxylin and eosin (H&E) staining was performed by the Research Histology Lab in the Pitt Biospecimen Core. DNA fragmentation was assessed using the Click-iT® Plus TUNEL Assay Kit (ThermoFisher Scientific) according to the manufacturer’s instructions.

**Glutathione measurement:** Total hepatic GSH was measured in liver homogenates using the Glutathione Colorimetric Detection Kit (Invitrogen, Carlsbad, CA) according to the manufacturer’s instructions.

**RNA sequencing and analysis:** Total RNA was extracted from snap-frozen liver tissue using TRIzol reagent (Life Technologies, Carlsbad, CA). For baseline (0 hour) samples, RNA was isolated from individual mice (n = 4 per genotype). For the 12 to 96 hour time points, RNA from three biological replicates per genotype per time point was pooled prior to sequencing (n = 3 mice pooled per group). All RNA samples were submitted to Novogene (Sacramento, CA) for quality control, library preparation, and sequencing. Adapter sequences and low-quality reads were removed using Trimmomatic (v0.38).^1^ Clean reads were aligned to the Mus musculus reference genome (mm10) using STAR aligner (v2.6.1a),^2^ and gene-level read counts were quantified with --quantMode GeneCounts option. Subsequently, differential gene expression was assessed using DESeq2, with a false discovery rate (FDR) of 0.05 and a minimum fold change of 1.5 used to define significantly differentially expressed genes. Canonical pathway and upstream regulator analyses were conducted using Ingenuity Pathway Analysis (IPA; Qiagen, version 01-10), based on differentially expressed genes and associated downstream expression signatures. In addition, tight clustering was performed to cluster genes with similar expression levels across different time points.^3^ The raw RNA-seq data were submitted to the Gene Expression Omnibus database (http://www.ncbi.nlm.nih.gov/geo; accession number GSE303454; token chodccqaprinxgh).

***In Vitro* Treatment of WT Hepatocytes with APAP:** Primary hepatocytes were isolated from adult (2-3 months old; male) and 3 million viable cells were seeded into 10-cm Primaria cell culture plates (Corning) in seeding media consisting of DMEM/F-12 with 15 mM HEPES (Corning), 5% fetal bovine serum (FBS; Atlanta Biologicals, Atlanta, GA), and Antibiotic-Antimycotic Solution (Corning). After 4 hours, the seeding medium was replaced with growth medium composed of DMEM/F-12 with 15 mM HEPES, 0.5% FBS, Antibiotic-Antimycotic Solution, and ITS Supplement (1 μg/mL insulin, 0.55 μg/mL transferrin, and 0.67 ng/mL sodium selenite; Gibco Life Technologies). Cells were allowed to adhere overnight. The following morning, cells were treated with 0-10 mM APAP diluted in growth medium. After 24 hours, cells were imaged, trypsinized, and stained with FVD780 and Hoechst for assessment of viability and ploidy (see Ploidy Analysis section).

**Microscopy:** Micrographs were captured with a TiU fluorescent microscope (Nikon, Melville, NY) equipped with a Moment sCMOS monochrome camera (fluorescent images) (Photometrics, Tuscon, AZ) or Nikon DS-Fi3 color camera (non-fluorescent images). Images were processed with Nikon NIS Elements Advanced Research software and QuPath v0.4.2.^4^

**Detailed information for select reagents**

| Primary Antibodies | | | |
| --- | --- | --- | --- |
| *Epitope* | *Antibody with species reactivity* | *Vendor* | *Catalog #* |
| PCNA | Mouse anti-PCNA | Cell Signaling | 2586 |
| β-ACTIN | Rabbit anti-β-ACTIN | Cell Signaling | 4970 |
| Cyclin D1 | Rabbit anti-Cyclin D1 | Cell Signaling | 55506 |
| β-CATENIN | Rabbit anti-β-CATENIN | Cell Signaling | 8480 |
| Phospho-β-CATENIN | Rabbit anti-Phospho-β-CATENIN | Cell Signaling | 9561 |
| Non-phospho β-CATENIN | Rabbit anti-Non-phospho β-CATENIN | Cell Signaling | 19807 |
| SAPK/JNK | Rabbit anti-SAPK/JNK | Cell Signaling | 9252 |
| Phospho-SAPK/JNK | Rabbit anti-Phospho-SAPK/JNK | Cell Signaling | 4668 |
| SIRT6 | Rabbit anti-SIRT6 | Cell Signaling | 12486 |
| SIRT1 | Rabbit anti-SIRT1 | Cell Signaling | 3931 |
| SIRT3 | Rabbit anti-SIRT3 | MilliporeSigma | 07-1596 |
| CYP2E1 | Rabbit anti-CYP2E1 | MilliporeSigma | ab1252 |
| CYP1A2 | Goat anti-CYP1A2 | Santa Cruz | sc-9835 |
| Secondary Antibodies | | | |
| *Species reactivity* | *Antibody with conjugate* | *Vendor* | *Catalog #* |
| Rabbit | Anti-rabbit IgG | Cell Signaling | 7074 |
| Mouse | Anti-mouse IgG | Cell Signaling | 7076 |

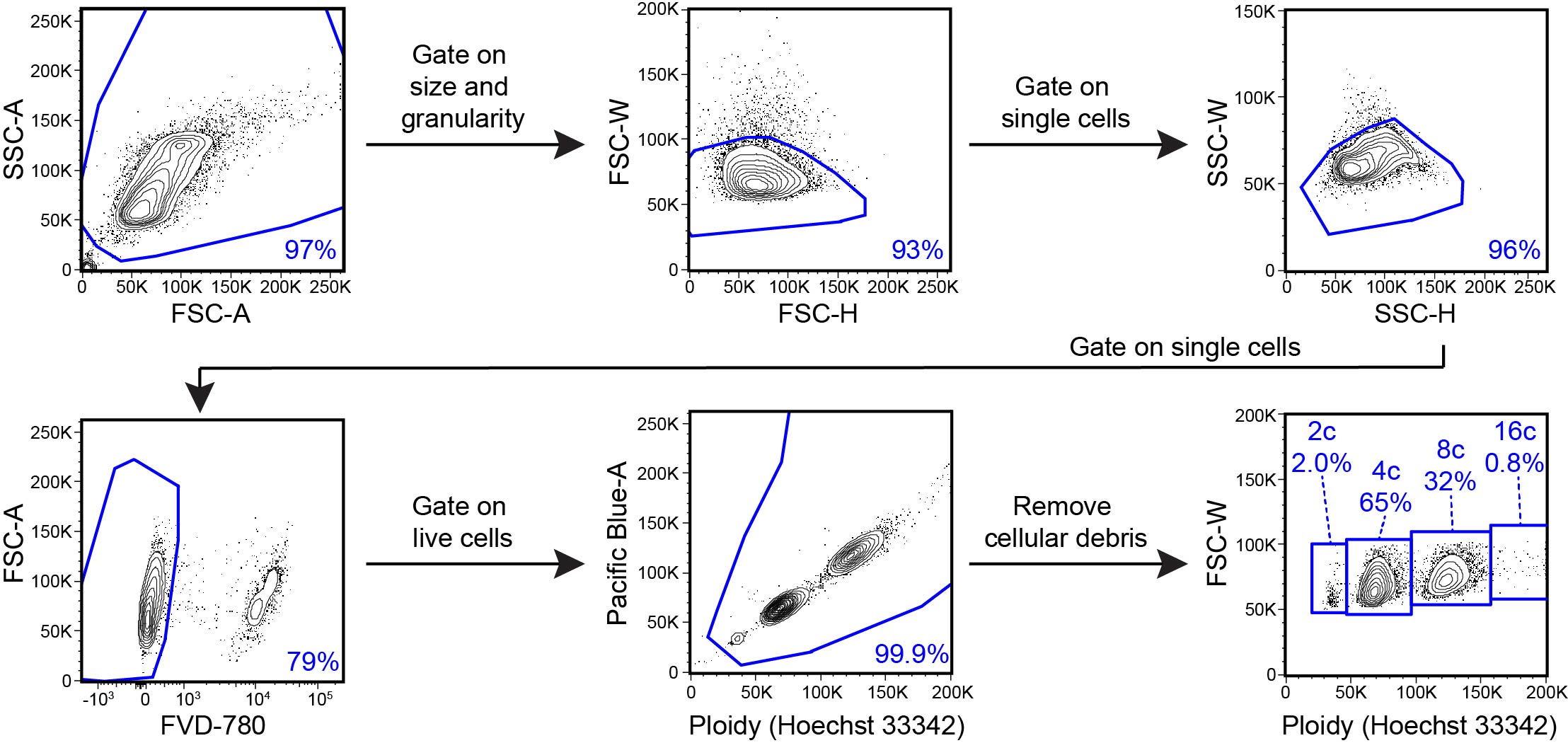

**Supplemental Figure S1. Flow cytometry gating strategy for analysis of hepatocyte ploidy distribution.**

Single-cell suspensions of hepatocytes from control and LKO mice were stained with FVD780 (fixable viability dye) and Hoechst. Cells were first gated based on size and granularity, then on single cells, followed by exclusion of dead cells and debris. Ploidy populations were determined by Hoechst 33342 fluorescence intensity. Shown plots are representative of the control mouse in Figure 1B.

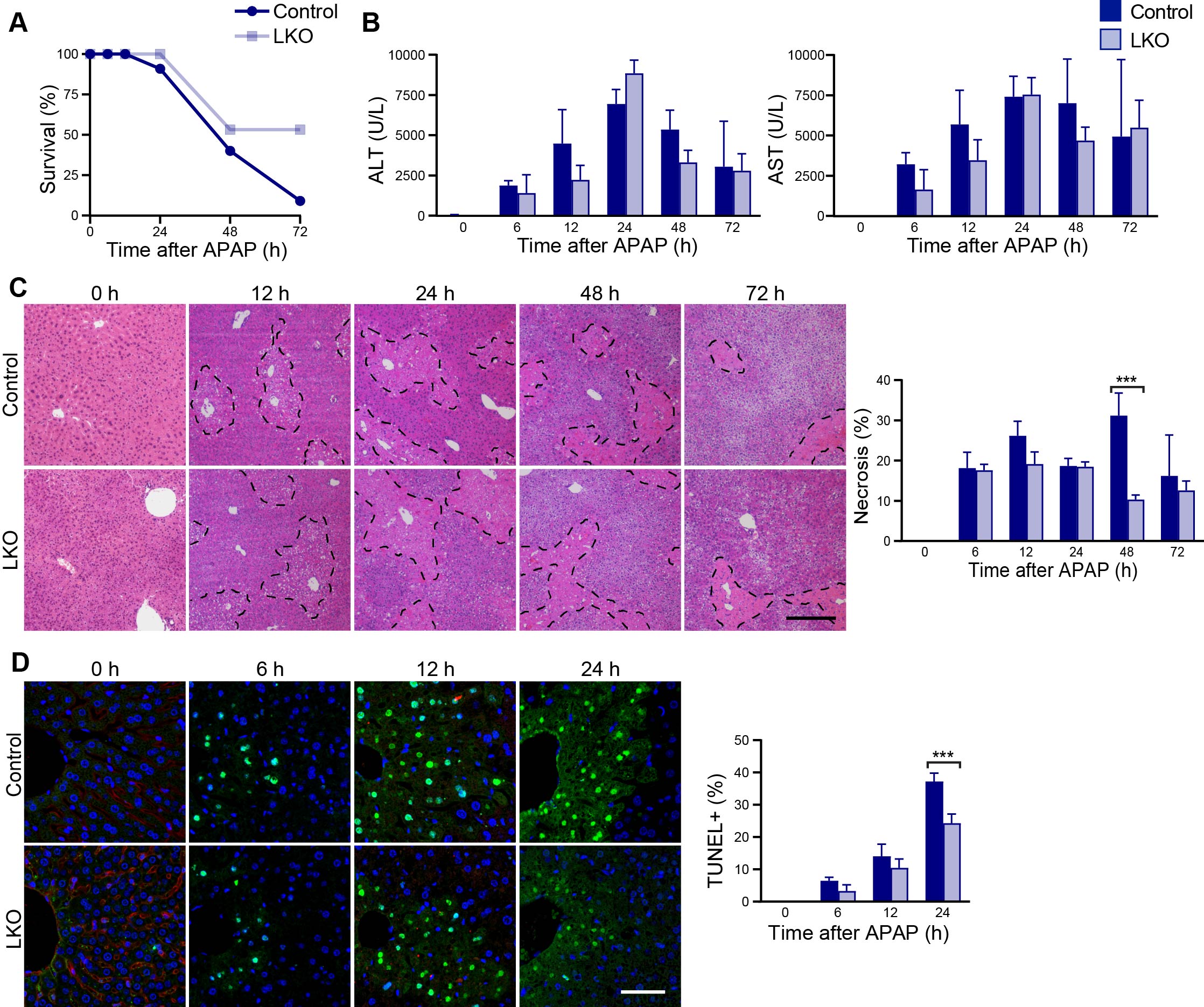

**Supplemental Figure S2. LKO mice resist lethal dose of APAP (600 mg/kg). (A)** Survival curves of control and LKO mice following 600 mg/kg APAP overdose (n = 4-12/genotype/timepoint). **(B)** Levels of liver biomarkers ALT and AST in the serum (n = 4-12/genotype/timepoint). **(C)** Quantification of necrosis by H&E Staining (n = 4-12/genotype/timepoint). Scale bar = 200 µm. **(D)** TUNEL staining showing DNA fragmentation (green) and nuclear staining with Hoechst 33342 (blue) (n = 4-9/genotype/timepoint). Scale bar = 50 µm. Representative images, plots, and quantification results are shown. Graphs show mean ± SEM. *P < 0.05; **P < 0.01; ***P < 0.001.

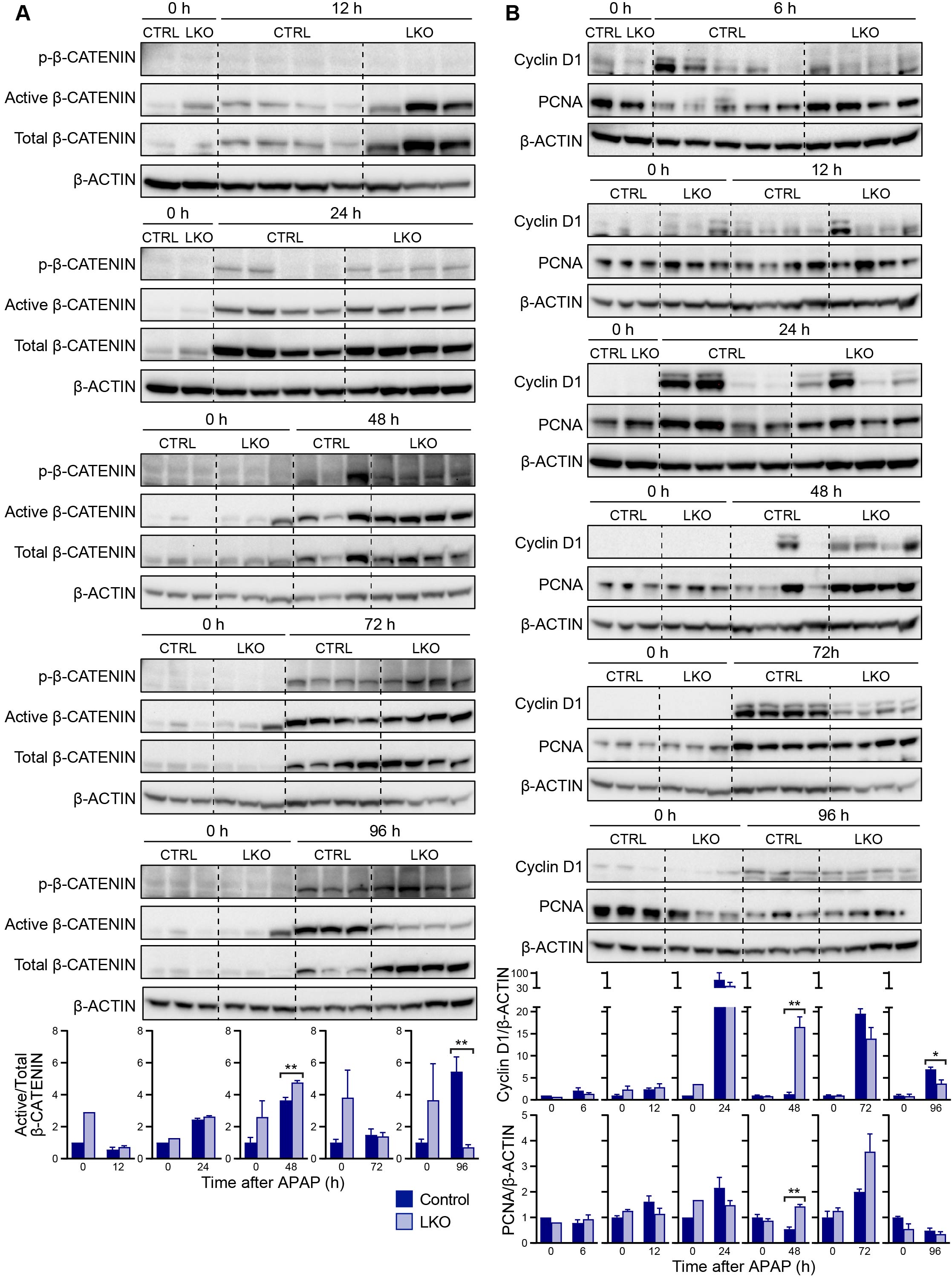

**Supplemental Figure S3. LKO mice show enhanced proliferation at 48 hours. (A-B)** Western blotting of whole liver lysates collected from control and LKO mice 0-96 hours after 300 mg/kg APAP overdose, showing expression of proteins involved in proliferation: **(A)** active β-CATENIN relative to total β-CATENIN (n = 3-4/genotype/timepoint), **(B)** Cyclin D1 and PCNA relative to β-ACTIN where one sample at 48 hours was identified as an outlier using the Grubbs Test with an alpha = 0.01 per treatment group (n = 3-5/genotype/timepoint). Shown are representative blots and quantification results normalized to the 0 hour control, which is set to 1. Graphs show mean ± SEM. *P < 0.05; **P < 0.01; ***P < 0.001.

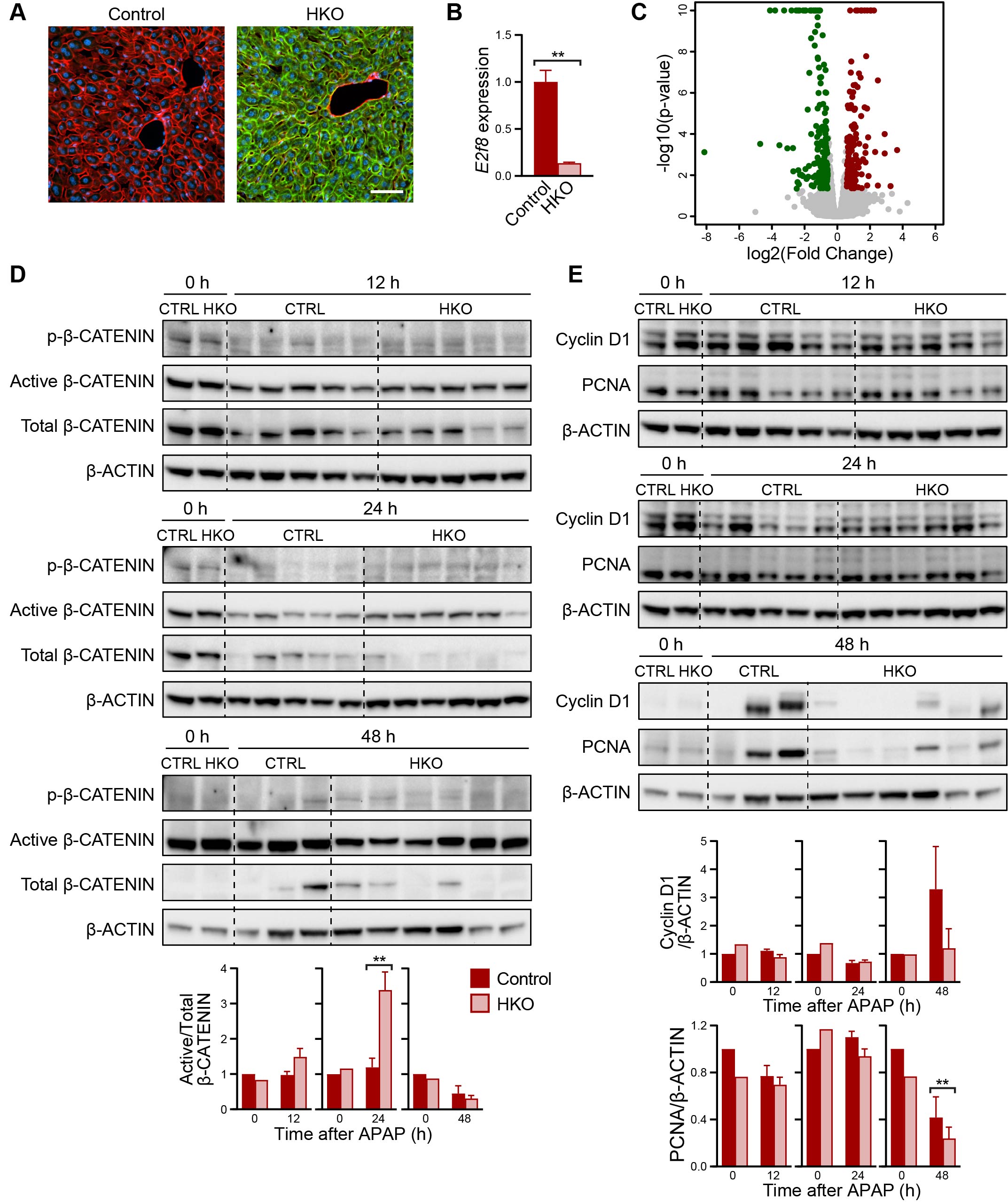

**Supplemental Figure S4. Expression in the HKO model. (A)** Representative fluorescent images of liver sections from control and HKO mice two weeks after AAV8 injection. In the R26R-mTmG Cre-reporter system, all cells express membrane-bound tdTomato (red) prior to Cre recombination. Upon Cre-mediated excision, tdTomato is replaced by membrane-bound GFP (green), indicating successful recombination. All hepatocytes in control livers remained tdTomato⁺, while >99% of hepatocytes in HKO livers were GFP⁺, demonstrating efficient, hepatocyte-specific Cre recombinase activity. Scale bar = 50 µm. **(B)** Expression of *E2f8* in control and HKO livers, based on RNA-seq gene counts in exons 3 and 4. Quantification is normalized to control, which is set to 1. **(C)** Volcano plot showing differential gene expression between control and HKO mice. Differentially expressed genes were defined by FDR = 5% and fold change ≥ 1.5. A total of 141 genes were upregulated (red) and 179 downregulated (green) in HKO compared to controls (Supplemental Table S1). **(D-E)** Western blotting of whole liver lysates collected from control and HKO mice 12, 24, and 48 hours after APAP overdose, showing expression of proteins involved in proliferation **(C)** active β-CATENIN relative to total β-CATENIN and **(D)** CCND1 and PCNA relative to β-ACTIN. Shown are representative blots and quantification results normalized to the 0 hour control, which is set to 1. Graphs show mean ± SEM. *P < 0.05; ** P < 0.01; ***P < 0.001.

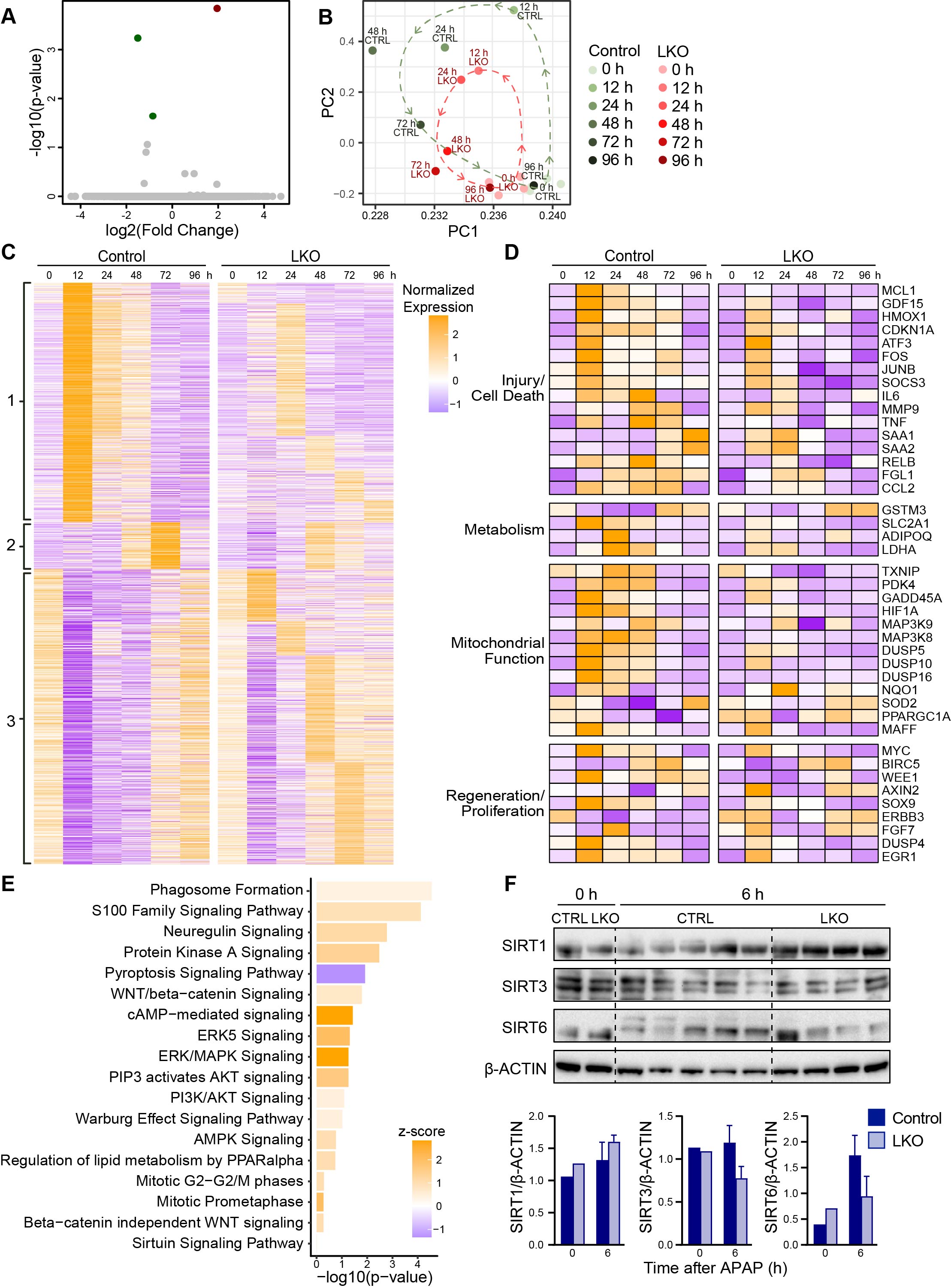

**Supplemental Figure S5. Gene and protein expression in LKO livers. (A)** Volcano plot showing differential gene expression between control and LKO mice at baseline (0 hours; n=4/genotype). Differentially expressed genes were defined by FDR = 5% and fold change ≥ 1.5. See Supplemental Data Table S2 for gene list. **(B)** PCA of RNA-seq data. Control samples (green) and LKO samples (red) are shown (0 hour n=4/genotype; 12-96 hour samples are pooled with n=3/genotype/timepoint). **(C)** Heat map showing expression patterns of differentially expressed genes grouped into clusters 1-3 based on temporal expression patterns. See Supplemental Table S3 for gene lists. **(D)** Heat map of differentially expressed genes categorized by functional pathways, including injury/cell death, metabolism, mitochondrial function, and regeneration/proliferation. See Supplemental Table S4 for gene lists. **(E)** IPA of control and LKO livers at 12 hours post-APAP (pooled samples; n=3/genotype). Relevant enriched pathways are shown. **(F)** Western blot analysis of SIRT1, SIRT3, and SIRT6 protein levels at 0 and 6 hours post-APAP treatment. Representative blots and quantification are shown. Data are normalized to control at baseline and expressed as mean ± SEM.

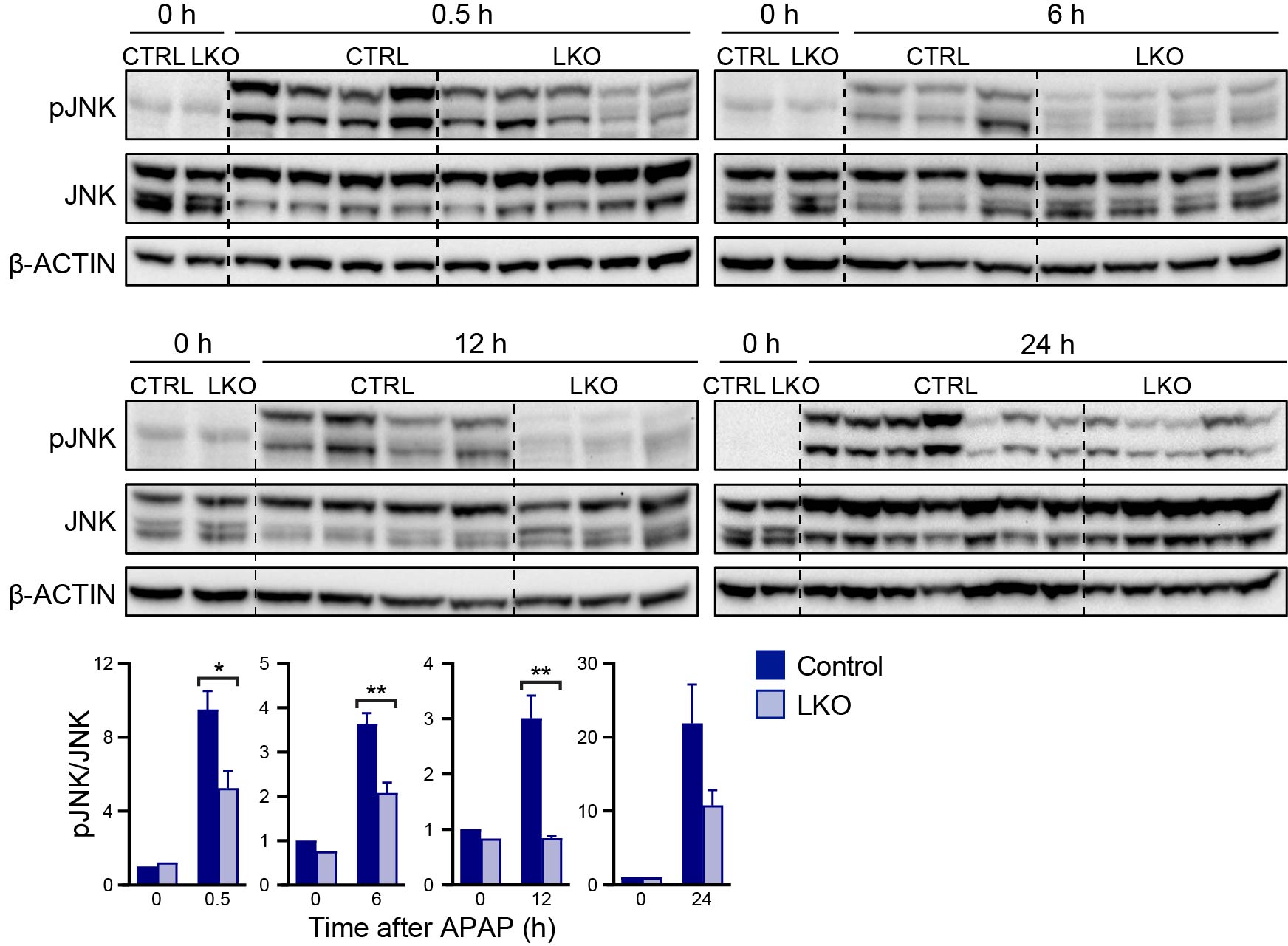

**Supplemental Figure S6. Reduced JNK activation in LKO mice after APAP overdose.** Western blots of pJNK relative to total JNK in whole liver lysates at 0-24 hours after APAP treatment in LKO and control mice (n = 3-6/genotype/timepoint). Representative blots and quantification are shown. Data are normalized to control at baseline and expressed as mean ± SEM. *P < 0.05; **P < 0.01; ***P < 0.001.
